# Yeast surface display-based identification of ACE2 mutations that modulate SARS-CoV-2 spike binding across multiple mammalian species

**DOI:** 10.1101/2021.03.16.435705

**Authors:** Pete Heinzelman, Jonathan C. Greenhalgh, Philip A. Romero

## Abstract

Understanding how SARS-CoV-2 interacts with different mammalian angiotensin-converting enzyme II (ACE2) cell entry receptors elucidates determinants of virus transmission and facilitates development of vaccines for humans and animals. Yeast display-based directed evolution identified conserved ACE2 mutations that increase spike binding across multiple species. Gln42Leu increased ACE2-spike binding for human and four of four other mammalian ACE2s; Leu79Ile had a effect for human and three of three mammalian ACE2s. These residues are highly represented, 83% for Gln42 and 56% for Leu79, among mammalian ACE2s. The above findings can be important in protecting humans and animals from existing and future SARS-CoV-2 variants.

## Introduction

The SARS-CoV-2 coronavirus has demonstrated substantial ability to mutate in ways that increase its transmissibility among humans [1], reduce the effectiveness of COVID-19 vaccines [2], and enable bidirectional transmission of the virus between humans and animals [3]. These observations have motivated the development of novel COVID-19 vaccines and vaccine boosters for humans [4], vaccination agents for economically important animals [5], prospecting efforts centered on isolating viruses from wild animals to identify new coronavirus strains that possess potential to be transmitted to the human population [6], and establishment of livestock husbandry and coronavirus screening practices to reduce the likelihood of coronavirus infections spreading within farmed animal populations [7].

The above pursuits demand a deeper understanding of how sequence variation in both the ACE2 receptor and the viral spike protein influences transmission within the human population and across species barriers. In this work we explore how mutations in human and other mammalian ACE2s affect spike protein binding using a yeast surface display-based screening method [8]. We discovered that substitutions at ACE2 amino acid positions 34, 42, and 79 increased spike binding in several ACE2 orthologs. The increased binding of Gln42Leu and Leu79Ile mutations are especially noteworthy given that Gln appears at position 42 in 83% of annotated mammalian ACE2s, while Leu is present at position 79 in 56% of such proteins [9].

The above finding that ACE2 mutations have similar effects upon spike binding across multiple ACE2 orthologs suggests that mutations in the spike protein can also modulate the ACE2-spike binding interaction in a species-independent manner. It follows that mutant SARS-CoV-2, or other emergent coronaviruses, could feature spike proteins that impart cross-species transmission propensities greater than those for SARS-CoV-2 strains observed to date, thus motivating proactivity and breadth of action with respect to vaccine development, novel coronavirus strain prospecting, and animal farming practices aimed at preventing and containing coronavirus infection. Combining our results with structural analysis of ACE2-spike binding interactions, comparisons of ACE2 sequences across mammalian species, and existing knowledge of SARS-CoV-2 mutations that increase spike binding *in vitro* [10] will provide a global view of viral emergence and transmission that will help mitigate the societal and economic impacts of current and future coronavirus outbreaks.

## Results

Our work focused on four key mammalian ACE2 orthologs from *Homo sapiens, Felis catus* (domestic cat), *Canis familiarus* (domestic dog), and *Sus scrofa* (domestic pig). We cloned genes encoding the peptidase domains (residues 19-614) of these four ACE2 orthologs into a yeast surface display vector (Supporting Figure 1) and measured their expression and spike binding using flow cytometry. As shown in Supporting Figure 2, all four of these ACE2s were well-displayed on the yeast surface and bound to recombinant SARS-2-CoV spike receptor binding domain (RBD) with the following rank order of flow cytometry assay binding signal values: human > cat ≈ pig > dog.

**Figure 1.**
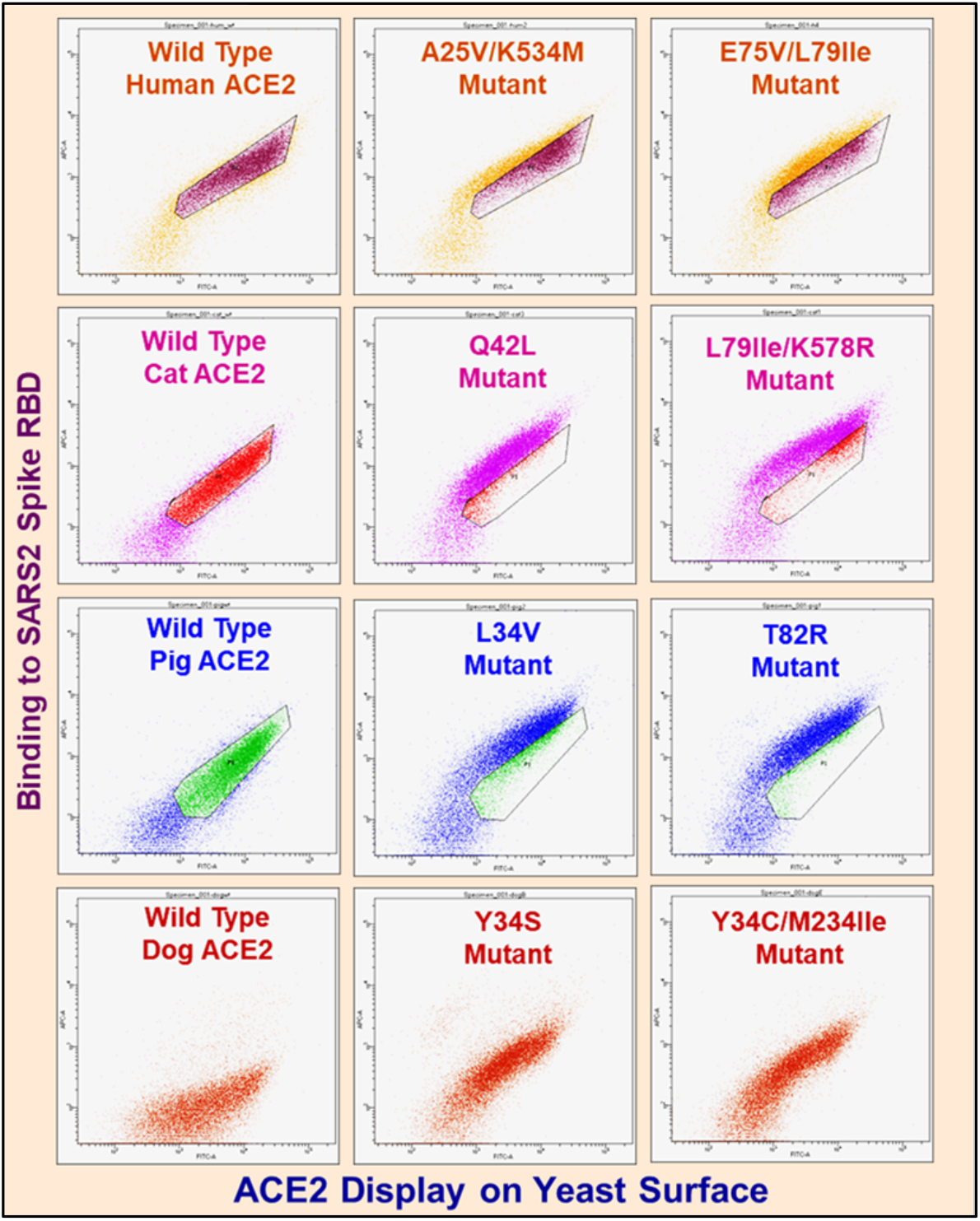
Flow cytometry dot plots for wild type and ACE2 random mutant library clones enriched after three rounds of FACS. X-axes denote Alexa488 fluorescence (ACE2 display). Y-axes denote Alexa647 fluorescence (ACE2 binding to spike protein). Plots depict dots for approximately 3*10^4^ yeast cells. ACE2 yeast incubated with spike as follows: human - 1 nM, cat - 4 nM, pig - 4 nM, dog - 25 nM. For poorly understood biological reasons homogeneous populations of yeast carrying identical display plasmids feature 25% or greater cells (lower left of plots) that do not display protein.

**Figure 2.**
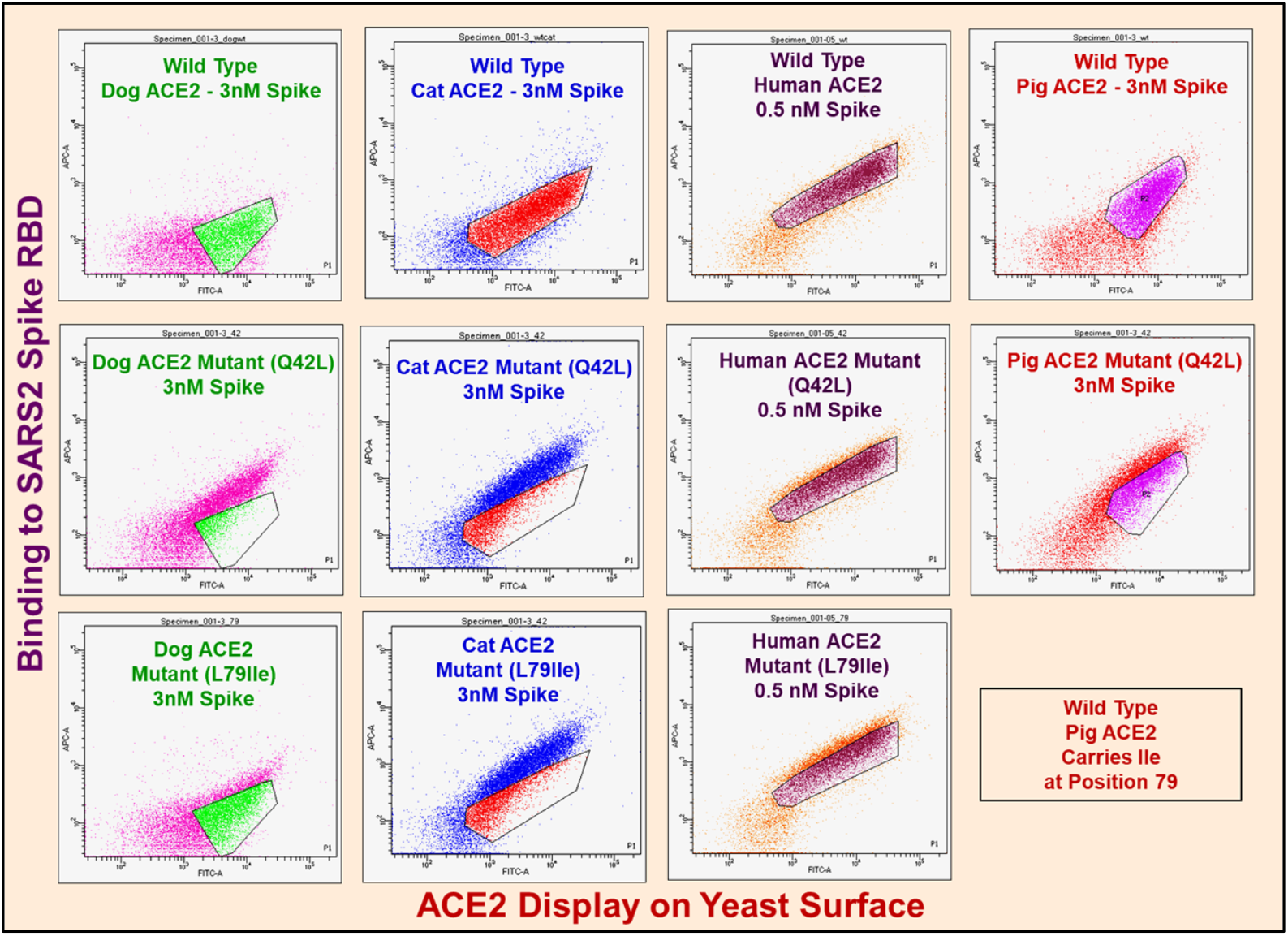
Dot plots for wild type ACE2 and Gln42Leu single mutants. X-axes denote Alexa488 fluorescence (ACE2 display). Y-axes denote Alexa647 fluorescence (binding to spike). Plots depict dots for approximately 3*10^4^ yeast cells. ACE2 yeast incubated with spike as denoted in figure.

We performed random mutagenesis on the four ACE2 orthologs [11] and used three rounds of FACS-based enrichment to isolate high affinity mutants for each ortholog. Yeast-displayed ACE2 libraries contained > 10^5^ transformants and sequenced clones featured 1-3 coding mutations per gene; these numbers ensured comprehensive screening of ACE2 mutational space.

After the third round of FACS we performed flow cytometric analysis and DNA sequencing to confirm that we had isolated multiple unique high affinity clones from each library (Figure 1 - preceding page & Supporting Table 1).

We focused our further analysis on mutations that enhance spike binding and that also occur at highly conserved sites across annotated mammalian ACE2s [9]. The fact that mutations could enhance binding suggests these sites were originally suboptimal for binding and may be primed for the evolving virus to exploit. We identified positions 42 (83% percent representation for Gln) and 79 (56% representation for Leu) because they were present in our high affinity clones and are conserved across annotated mammalian ACE2s. Although ACE2s carrying mutations at position 34 were enriched in the cat, pig, and dog sorts this position is relatively variable, with His being most highly conserved at 34%, and thus not an ideal fit to the objective of assessing whether identical substitutions can increase spike binding affinity across multiple ACE2 orthologs.

We individually introduced the Gln42Leu and Leu79Ile mutations, where it is notable that among annotated farmed animal ACE2s Ile79 in pig is the only instance of identity with these substitutions, into each of the mammalian ACE2s and found both of these mutations substantially increased spike-binding signals (Figure 2).

The Gln42Leu mutation increased the binding signal 10-400% across the four ACE2 orthologs, while Leu79Ile’s increases were in the 30-90% range.

Following on this result, we sought to further explore whether the Gln42Leu mutation would enhance spike binding in other mammalian ACE2s. We identified the ACE2 from the halcyon bat (*Rhinolophus alcyone*) as an additional ortholog that carries Gln at position 42, is compatible with our yeast display system, and binds to spike RBD (Supporting Figure 3). We found the Gln42Leu mutation also enhanced spike binding of the halcyon bat ACE2 (Supporting Figure 4). Finally, we tested the effect of the Gln42Leu in all five ACE2s across a wide range of spike RBD concentrations (Figure 3).

**Figure 3.**
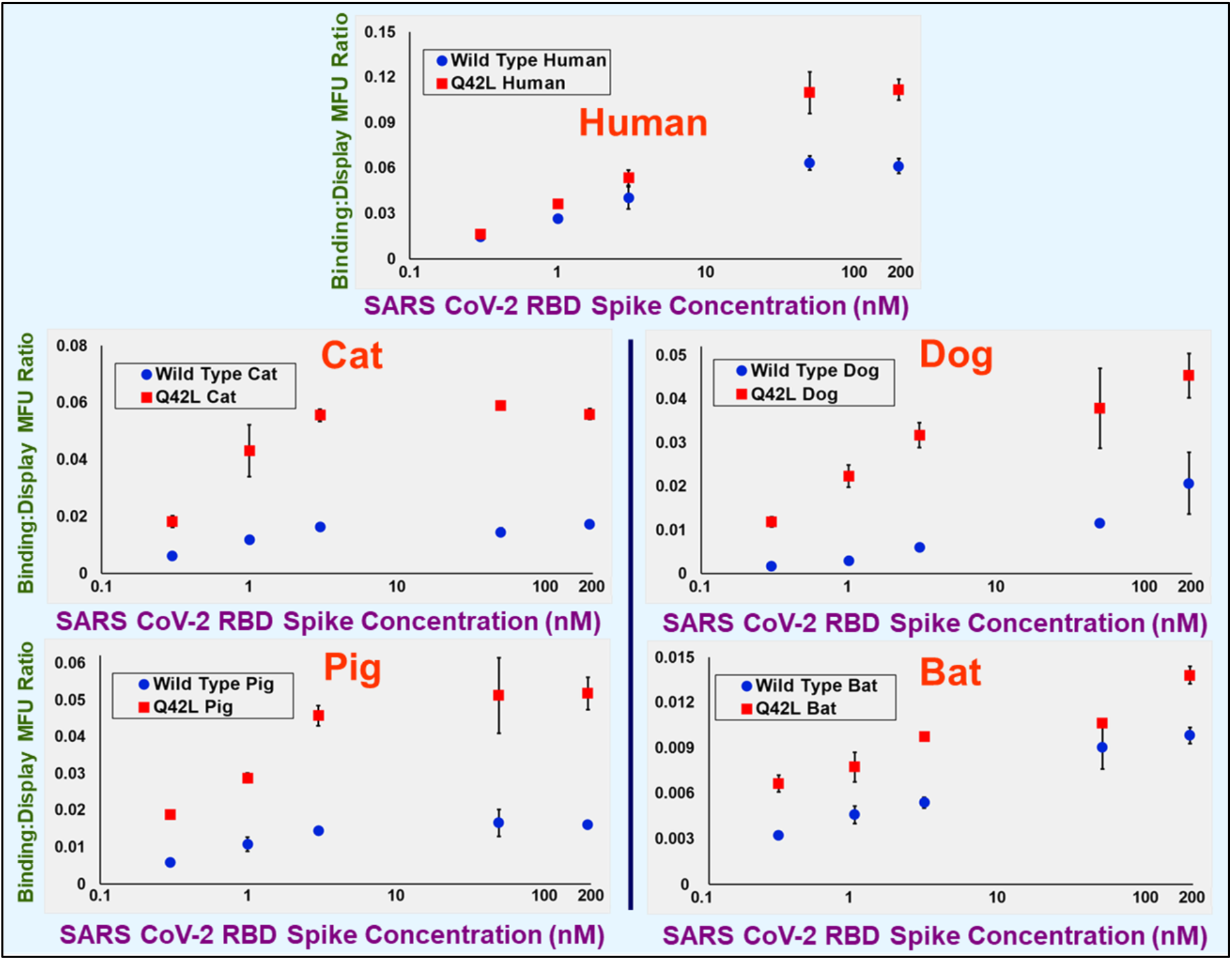
Normalized yeast display flow cytometry spike binding assay MFU values versus spike RBD concentration for wild type and Gln42Leu mutant ACE2s. Normalized values in plots are spike binding MFU divided by ACE2 display MFU. Binding MFU (Alexa647) are MFU values for gated, *myc*-positive ACE2-displaying yeast populations minus MFU for gated, *myc*-negative nondisplaying yeast. Display MFU (Alexa488) are MFU values for gated, *myc*-positive ACE2-displaying yeast populations minus MFU for gated, *myc*-negative nondisplaying yeast. Values shown are mean of duplicate measurements.

**Figure 4.**
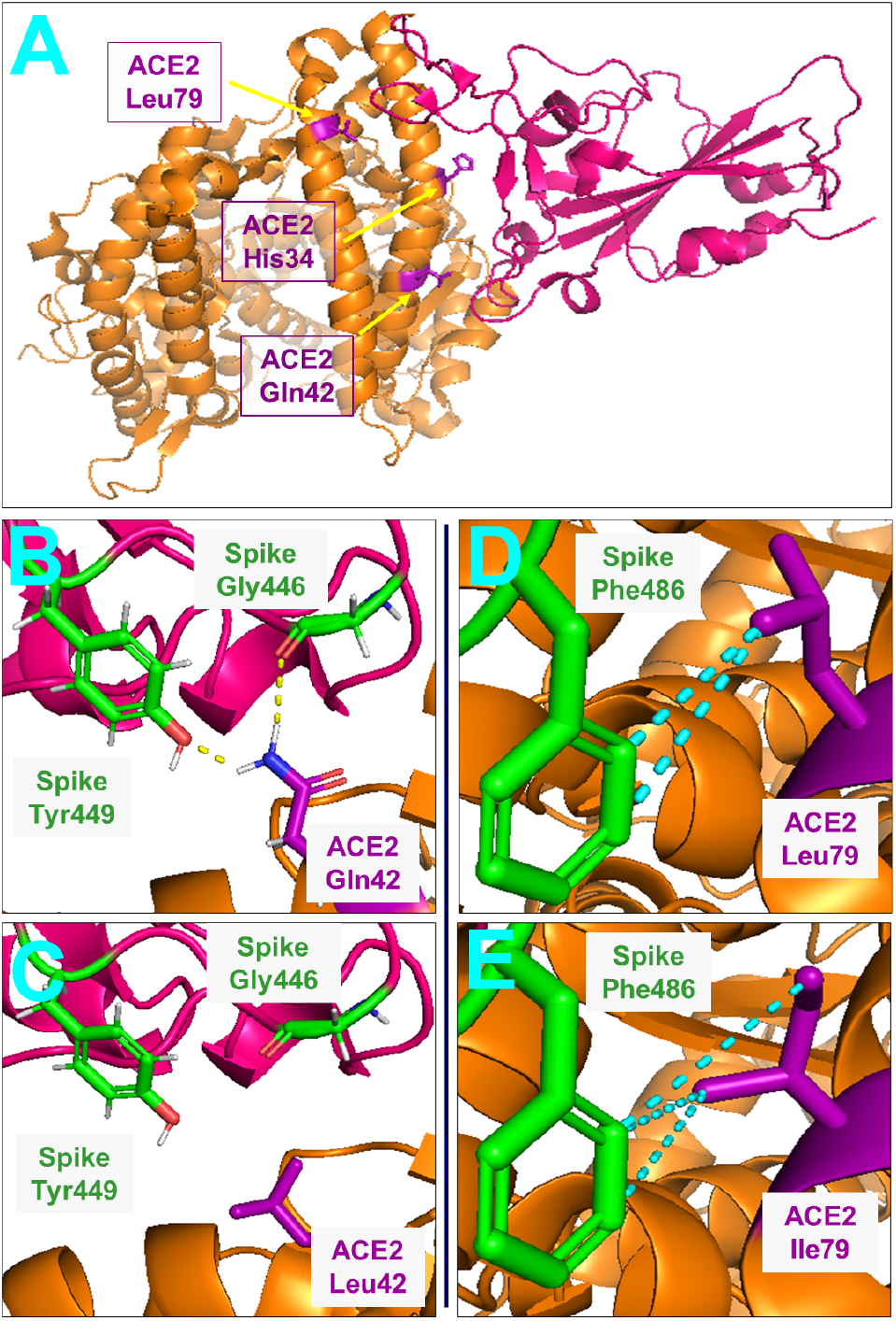
Structural views of SARS-CoV-2 spike-ACE2 complex. ACE2 in orange, spike RBD in pink. Panel A: Overall spike-ACE2 complex with ACE2 affinity-determinant positions identified by yeast display screening in purple. Panel B: Illustration of hydrogen bonds (yellow) formed between ACE2 Gln42 and spike Gly446 and Tyr449. Hydrogen atoms: white, oxygen: red, nitrogen: blue. Panel C: Absence of these hydrogen bonds for ACE2 Gln42Leu mutant. Panel D: Hydrophobic contacts (less than 5 Å carbon atom separation - colored in cyan) between ACE2 Leu79 and spike Phe486. Panel E: Hydrophobic contacts between spike Phe486 and ACE2 Leu79Ile mutant. Figures generated from Protein Data Bank structure 6m0j (human ACE2) using Pymol.

A competition, likely due to steric interference, between the primary and secondary antibodies used for *myc* tag detection and the respective spike RBD and fluorescent anti-His_6_ antibody used for spike detection results in the wild type and Q42L mutant ACE2s appearing to have similar half maximal spike binding concentrations. Given however, the increase in maximum normalized binding signal for all five mutant ACE2s it is clear that the Q42L substitution increases ACE2 spike binding affinity.

Rosetta [12] human ACE2-spike binding energy calculations returned ΔΔG values of 0.12 +/- 0.53 and 0.26 +/- 0.55 Rosetta Energy Units (REU) for respective Gln42Leu and Leu79Ile mutations. Although positive, these values are small and thus do not contradict the increased spike binding affinity observed in yeast display assays.

Analysis of the human ACE2-spike RBD co-crystal structure (pdb 6m0j) [13] shows that the Gln42Leu substitution eliminates hydrogen bonds between Gln42 and spike residues Gly446 and Tyr449 (Figure 4 - preceding page). Examination of Rosetta energy terms reveals that change in solvation energy (Δfa_sol) is dominant (−0.89 +/- 0.73 REU) and indicates that loss of these hydrogen bonds is overcome by favorable solvation effects at other ACE2 and spike residues.

Figure 4 illustrates the plausibility of increased hydrophobic contact with spike residue Phe486 contributing to the Leu79Ile ACE2 mutants’ increased spike binding affinity. Rosetta energy term analysis did not identify factors that may underlie the effect of this mutation.

## Discussion

In this work we performed directed evolution on four mammalian ACE2 orthologs to identify mutations that enhance binding to the SARS-CoV-2 spike protein. The identified clones revealed numerous and varied mechanisms to increase spike binding, highlighting the latent potential of SARS-CoV-2 to evolve a broad host range. We focused further analysis on ACE2 positions 42 and 79 because they are highly conserved across mammalian ACE2 homologs and mutations at these sites can enhance spike binding. We found the mutation Leu79Ile enhanced spike binding in 3/3 mammalian ACE2’s tested and mutation Gln42Leu enhanced spike binding in 5/5 ACE2s tested.

Our finding that ACE2 positions 34, 42 and 79, all of which reside within ACE2’s N-terminal helices, can play important roles in the ACE2-spike RBD binding interaction across multiple mammalian orthologs is in accord with prior studies of human ACE2. Procko and colleagues [14] observed that Gln42Leu and Leu79Ile mutations, as well as substitutions at His34, increased human ACE2’s binding affinity toward the spike RBD. Additionally, Wells *et al*. [15] identified mutations at residues His34 (His34Ala, His34Ser, and His34Val) and Leu79 (Leu79Pro) that increased human ACE2 spike RBD binding affinity.

The observed impact of Gln42Leu and Leu79Ile mutations on spike RBD binding interaction for multiple ACE2 orthologs may be leveraged in the context of crystal structure-guided computational prediction of SARS-CoV-2 spike RBD mutations that could give rise to new virus strains possessing increased propensity for cross-species transmission. Such predictive efforts can find a foundation in prior respective *in silico* studies aimed at elucidating how animal ACE2 amino acid sequence impacts SARS-CoV-2 spike binding [16] and identifying human ACE2 residues [17] that increase the strength of the ACE2-spike interaction. Additionally, our discovery of the conserved effects of Glu42Leu and Leu79Ile ACE2 mutations could be put to use in cell culture studies focused on resolving the ACE2 and spike RBD amino acid sequence determinants of SARS-CoV-2 infectivity for both humans and animals; Farzan and co-workers [18] have previously quantified the *in vitro* infection susceptibilities, where we note that spike-host cell binding affinity is one of mulitple factors that impact coronavirus infectivity [19], of respective human cell lines transfected with more than a dozen mammalian ACE2 orthologs upon challenge with both SARS-CoV-2 and related coronaviruses.

The findings presented in this work, as well as other related computational and experimental pursuits, will contribute to our understanding of how ACE2 and spike RBD amino acid sequences influence SARS-CoV-2 transmission among humans and other mammals. This heightened understanding could play an important role in developing novel vaccines for both humans and animals and contribute to global food security by facilitating the establishment of effective practices for ensuring the health of farmed animal populations.

### Experimental Procedures

#### ACE2 library generation and screening

Residues 18-615 of human (UniProt Q9BYF1), cat (Q56H28), dog (E2RR65) and pig (K7GLM4) ACE2 genes were synthesized as yeast codon-optimized gBlocks (Integrated DNA Technologies, Coralville, IA) and ligated NheI-MluI into yeast display vector VLRB.2D-aga2 (provided by Dane Wittrup, MIT); this vector fuses the aga2 protein to the ACE2 C-terminus (Supporting Figure 1). ACE2 genes contained His to Asn mutations at positions 376 and 380 to abolish ACE2 proteolytic activity. The GeneMorph II Kit (Agilent Technologies, Santa Clara, CA) was employed to generate ACE2 random mutant libraries using wild type ACE2 display plasmid as template and forward and reverse primers CDspLt (5’-GTCTTGTTGGCTATCTTCGCTG-3’) and CDspRt (5’-GTCGTTGACAAAGAGTACG-3’). Error prone PCR products from mutagenesis reactions were digested NheI to MluI and ligated into VLRB.2D-aga2. Ligation products were concentrated using the Zymoclean Clean & Concentrator 5 kit (Zymo Research, Orange, CA) and electroporated into 10G Supreme *E. coli* (Lucigen, Middleton, WI). Transformants were pooled, cultured in LB media containing 100ug/mL carbenecillin overnight at 30°C and plasmids harvested using the Qiagen (Valencia, CA) Spin Miniprep kit.

Yeast display *Saccharomyces cerevisiae* strain EBY100 was made competent using the Sigma-Aldrich (St. Louis, MO) transformation kit. Transformants were pooled, cultured in low-pH Sabouraud Dextrose Casamino Acid media (SDCAA, per liter - 20 g dextrose, 6.7 grams yeast nitrogen base (VWR Scientific, Radnor, PA), 5 g Casamino Acids (VWR), Citrate buffer (pH 4.5) - 10.4 g sodium citrate / 7.4 g citric acid monohydrate) at 30°C and 250 rpm for two days. For ACE2 mutant library display induction, a 5 mL Sabouraud Galactose Casamino Acid (SGCAA, per liter - Phosphate buffer (pH 7.4) - 8.6 g NaH2PO*H2O / 5.4 g Na2HPO4, 20 g galactose, 6.7 g yeast nitrogen base, 5 g Casamino Acids) culture was started at optical density, measured at 600 nm, of 0.5 and shaken overnight at 250 rpm and 20°C.

After induction 3*10^6^ yeast cells were harvested by centrifugation, washed in pH 7.4 Phosphate Buffered Saline (PBS) containing 0.2% (w/v) bovine serum albumin (BSA) and incubated overnight at 4°C with rotation at 18 rpm in 1.2 mL of PBS/0.2% BSA containing 3 μg/mL anti-*myc* IgY (Aves Labs, Tigard, OR) and various concentrations of His_6_-tagged SARS-CoV-2 spike RBD (Sino Biological, Chesterbrook, PA): 1.5nM (human), 5nM (cat), 5nM (pig) and 50nM (dog). These spike concentrations were used in all rounds of sorting. Following overnight incubation yeast were washed in PBS/0.2% BSA and rotated at 18 rpm in same buffer containing 5 μg/mL mouse anti-His6 IgG (BioLegend, San Diego, CA) conjugated with Alexa647 using N-hydroxysuccinimidyl ester chemistry (Molecular Probes, Eugene, OR) and 2 μg/mL Alexa488-conjugated goat anti-chicken IgG (Jackson ImmunoResearch, West Grove, PA) for one hour at 4°C. Yeast cells were washed and resuspended in ice cold PBS for sorting on a FACS Aria III (Becton Dickinson, Franklin Lakes, NJ) located in the UW-Madison Flow Cytometry Laboratory. Sorting gates were set to isolate ACE2 mutant library members at or above the 97th percentile with respect to spike RBD binding.

Isolated yeast were cultured overnight in low pH SDCAA media at 30°C with shaking at 250 rpm. The following morning 1*10^6^ yeast were harvested and induced as above for a second round of incubation with spike and sorting; this process was repeated after sort two in a third and final round of sorting. Yeast enriched in round three sorts were plated on synthetic dropout (SD) -Trp (MP Biomedicals, Irvine, CA) plates and grown at 30°C for two days.

#### Flow cytometric analysis and sequencing of ACE2 library clones enriched during FACS

Along with wild type ACE2 yeast, five colonies for each animal ACE2 library sort were picked into 4 mL of low pH SDCAA and grown overnight at 30°C with shaking at 250 rpm. Cultures were induced in 5 mL of SGCAA overnight at 20°C with shaking at 250 rpm; induction starting OD was 0.5. After induction yeast were harvested by centrifugation and washed as above. 2*10^5^ yeast were tumbled overnight at 4°C in 500 μL of PBS/0.2% BSA containing 3 μg/mL anti-*myc* IgY and various concentrations of His6-tagged spike RBD (Figure 2). Secondary labeling, washing, and resuspension in PBS were performed as above prior to analysis using a Fortessa (Becton Dickinson) flow cytometer in the UW-Madison Biochemical Sciences Building.

ACE2 library clones with greater than wild type binding affinity were repicked and recultured from the post-sort plates and plasmids rescued using the ZymoPrep Yeast Plasmid Miniprep II kit (Zymo Research). ACE2 genes were amplified using primers CDspLt and CDspRt, PCR reactions cleaned up using the Qiaquick PCR purification kit, and PCR products sequenced using the CDspLt and CDspRt primers.

#### Flow cytometric analysis of ACE2 Gln42Leu mutants

For the four above and greater halcyon bat (UniProt A0A0N7IQX6) ACE2s, Gln42Leu mutant genes were constructed by overlap extension PCR using wild type ACE2 gBlocks as template, digested with NheI and MluI, and ligated into VLRB.2D-aga2. Plasmids for Gln42Leu mutant and wild type ACE2s were transformed into EBY100 yeast made competent using the Zymo Research Frozen EZ Yeast Transformation II kit with transformants plated onto SD -Trp plates and grown at 30°C. As described above, single colonies were cultured, induced, labeled with spike RBD (Figure 3), and analyzed using the Fortessa flow cytometer.

#### Rosetta binding energy calculations

Thirty-five models for respective Gln42Leu and Leu79Ile mutations were constructed using the pdb 6m0j human ACE2-spike RBD co-crystal structure. Structures were optimized with 35,000 backrub steps using the flex_ddg package [12]. ΔΔG values were determined by subtracting ΔG for the wild type complex from ΔG for mutant complexes; Δfa_sol values were identically calculated. Reported ΔΔG and Δfa_sol values and uncertainties are means and standard deviations for the thirty-five models.

## Abbreviations & Nomenclature

ACE2: Angiotensin-Converting Enzyme II
fa_sol: Rosetta solvation energy
MFU: Mean Fluorescence Units
REU: Rosetta Energy Units
SD: Synthetic Dropout
SDCAA: Sabouraud Dextrose Casamino Acid Media
SGCAA: Sabouraud Galactose Casamino Acid Media

## Data availability

All data is contained within the manuscript.

## Author contributions

P.H. and P.A.R conceived the project. P.H. and J.C.G. performed the experiments. P.H., J.C.G. and P.A.R analyzed the data. P.H. and P.A.R. wrote the manuscript.

## Funding and additional information

This work was supported by the US National Institutes of Health (5R35GM119854). “The content is solely the responsibility of the authors and does not necessarily represent the official views of the National Institutes of Health.”

