## Supplemental information for "Yeast surface display-based identification of ACE2 mutations that modulate SARS-CoV-2 spike binding across multiple mammalian species"

File contains Supporting Figures S1-S4 and Supporting Table S1

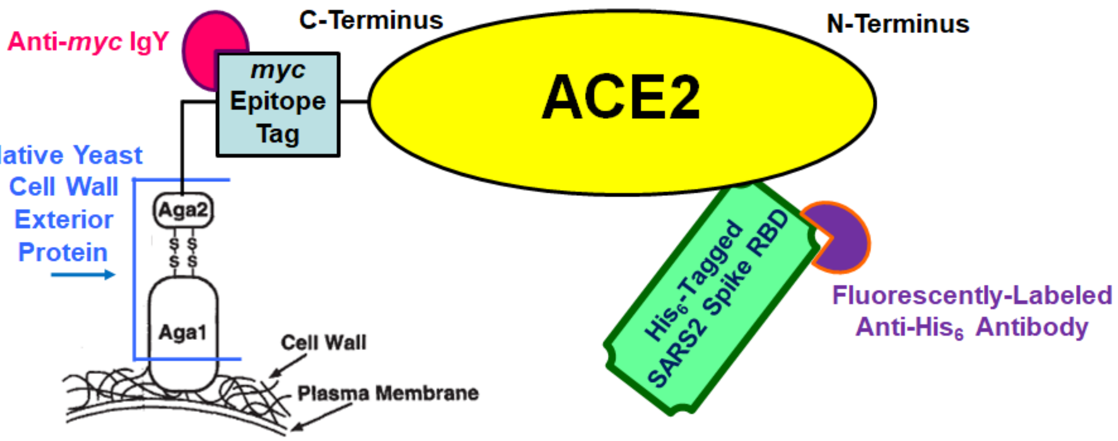

**Figure S1.** Yeast display schematic. ACE2 C-terminus linked to Aga2 native yeast surface protein on yeast cell wall. Anti-*myc* IgY (from chicken) used to quantify ACE2 display. Anti-myc IgY is not fluorescently labeled; fluorescent detection of *myc* epitope tag achieved via incubation with Alexa488-conjugated anti-chicken IgG (from goat) as secondary label (not depicted). Binding of yeast-displayed ACE2 to $His_6$-tagged SARS-CoV-2 spike RBD detected via incubation with Alexa647-conjugated anti-$His_6$ IgG (from mouse). Both ACE2 display and binding to spike RBD are measured by flow cytometry. Each yeast cell displays up to $10^4$ copies of a single ACE2 variant on its surface. Figure adapted from Boder and Wittrup (1997) *Nat Biotechnol.* 15:553-557.

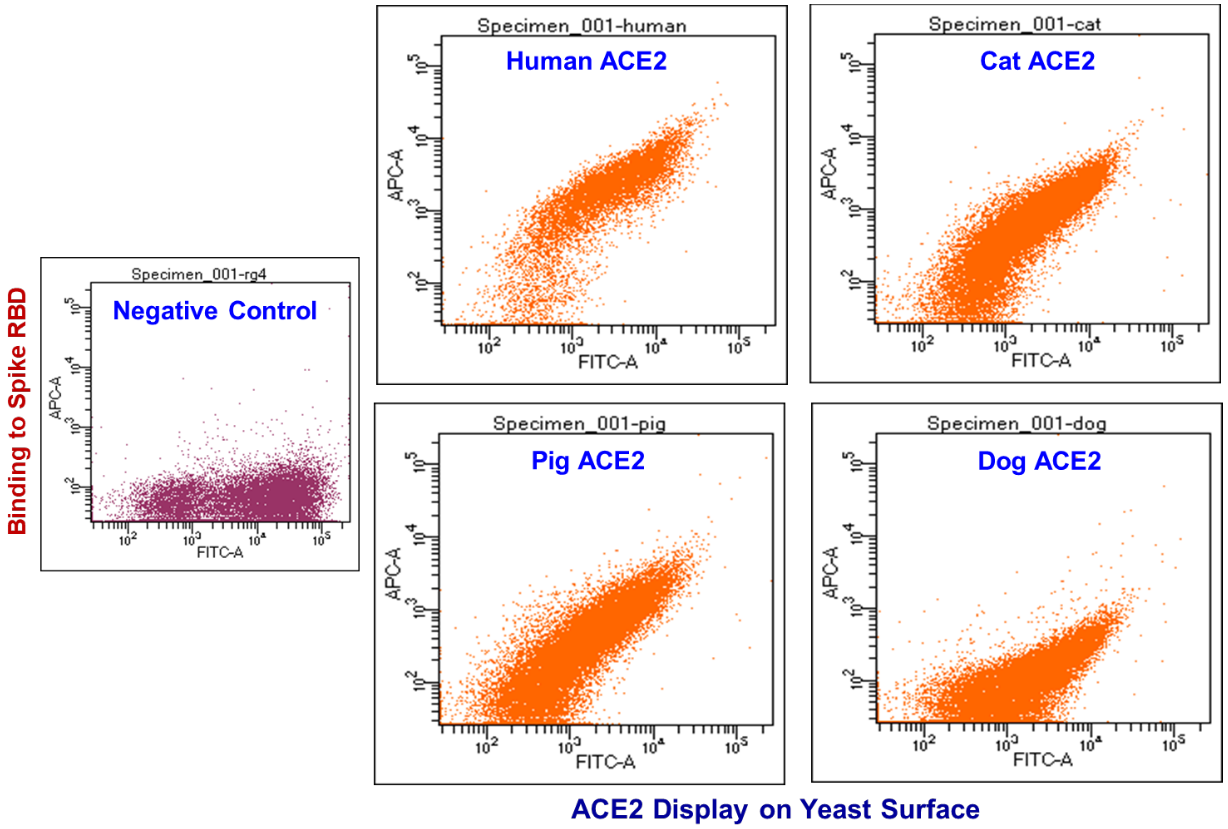

**Figure S2.** Flow cytometry dot plots for wild type human and other mammalian ACE2s. X-axes denote Alexa488 fluorescence (ACE2 display). Y-axes denote Alexa647 fluorescence (ACE2 binding to spike protein). Plots depict dots for approximately $2*10^4$ yeast cells. ACE2 yeast incubated with 250nM spike. For poorly understood biological reasons homogeneous populations of yeast carrying identical display plasmids feature 25% or greater cells (lower left of plots) that do not display protein. Negative control yeast carry display plasmid for horseradish peroxidase, a protein that displays on yeast but does not bind spike RBD.

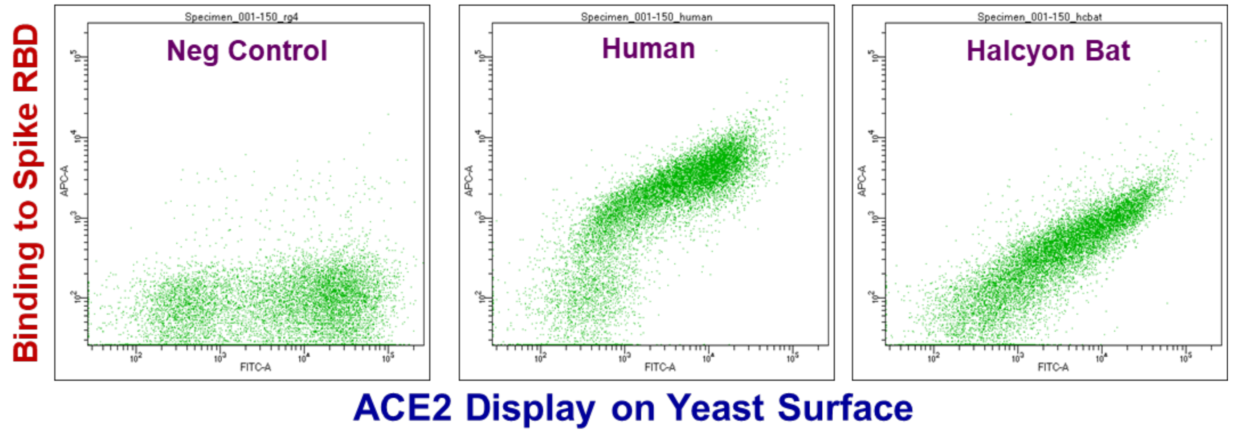

**Figure S3.** Flow cytometry dot plot for additional wild type mammalian ACE2, halcyon bat, that features Gln at position 42. X-axes denote Alexa488 fluorescence (ACE2 display). Y-axes denote Alexa647 fluorescence (ACE2 binding to spike protein). Plots depict dots for approximately $2*10^4$ yeast cells. ACE2 yeast incubated with 150nM spike RBD. For poorly understood biological reasons homogeneous populations of yeast carrying identical display plasmids feature 25% or greater cells (lower left of plots) that do not display protein. Negative control yeast carry display plasmid for horseradish peroxidase, a protein that displays on yeast but does not bind spike RBD.

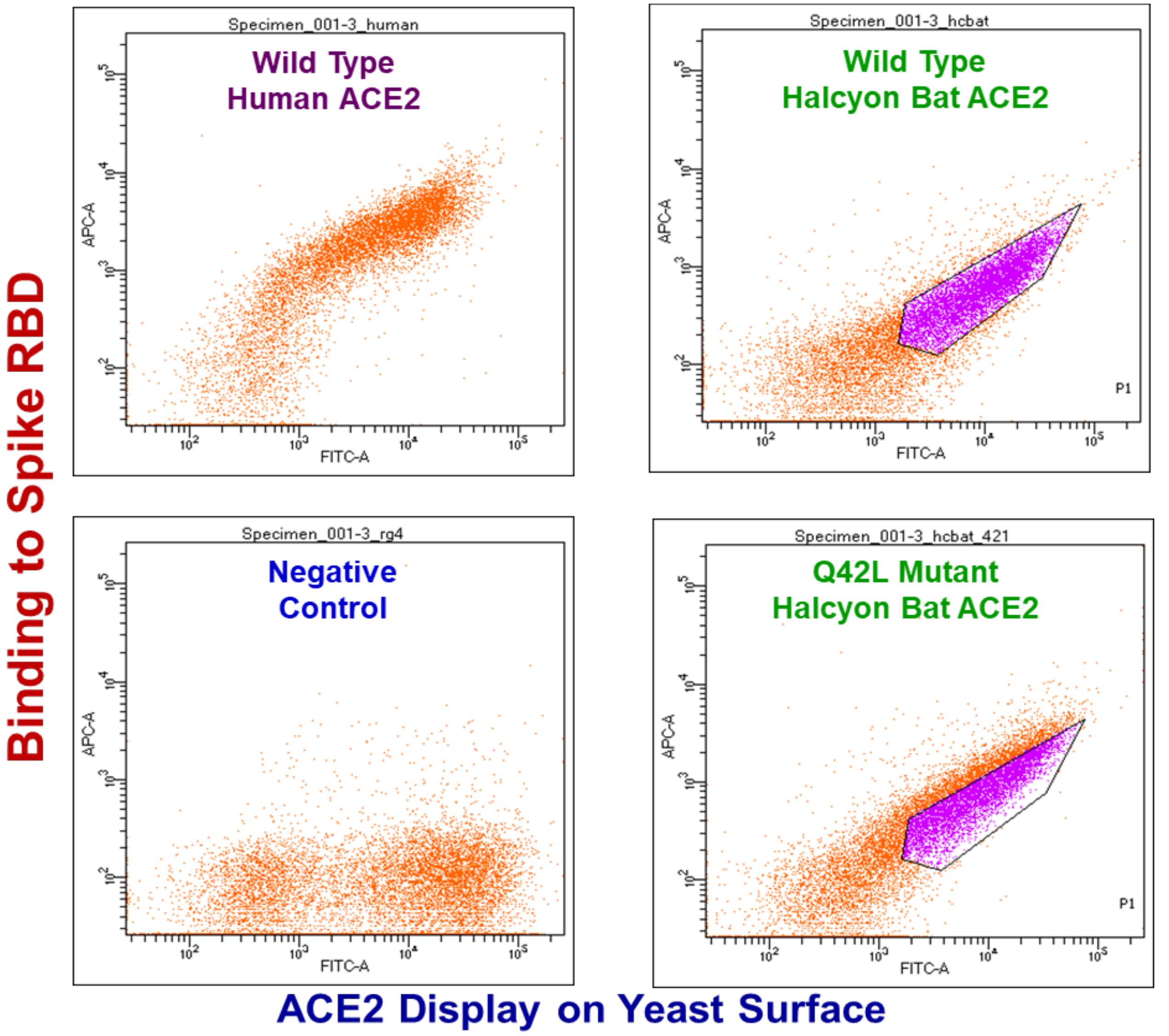

**Figure S4.** Dot plots for wild type halcyon bat ACE2 and Gln42Leu single mutant. X-axes denote Alexa488 fluorescence (ACE2 display). Y-axes denote Alexa647 fluorescence (binding to spike). Plots depict dots for approximately $3*10^4$ yeast cells. ACE2 yeast incubated with 3 nM spike RBD. For poorly understood biological reasons homogeneous populations of yeast carrying identical display plasmids feature 25% or greater cells (lower left of plots) that do not display protein. Negative control yeast carry display plasmid for horseradish peroxidase, a protein that displays on yeast but does not bind spike RBD.

**Table S1.** Mutations identified in high affinity ACE2 mutants isolated after three rounds of FACS for human, cat, dog and pig libraries. Table includes sequences for ACE2 mutant clones with higher than wild type spike binding signals determined by flow cytometric analysis as described in Experimental Procedures.

| Species | Clone Number | Amino Acid Substitutions |
|---|---|---|
| *Homo sapiens* (Human) | One | Q81R, K363N, E406Q |
| *Homo sapiens* (Human) | Two | A25V, K534M |
| *Homo sapiens* (Human) | Four | E75V, L79Ile |
| *Felis catus* (Cat) | Two | H34P |
| *Felis catus* (Cat) | Three | L79Ile, N90S, D213N |
| *Felis catus* (Cat) | Four | Q42L |
| *Felis catus* (Cat) | Five | L79Ile, K578R |
| *Sus scrofa* (Pig) | One | T82R |
| *Sus scrofa* (Pig) | Two | L34V |
| *Sus scrofa* (Pig) | Three | T82A, G215R, H239Q |
| *Sus scrofa* (Pig) | Four | L34V |
| *Canis familiaris* (Dog) | One | Y34S |
| *Canis familiaris* (Dog) | Two | Y34S |
| *Canis familiaris* (Dog) | Five | Y34C, M233Ile |
